# Prolactin Receptor Long Isoform ΔS1 Expression is Reduced in the Dorsal Root Ganglia of Humans with Chronic Pain

**DOI:** 10.64898/2026.10.27.762326

**Authors:** Lucy He, Keerthana Natarajan, Stephanie Shiers, Asta Arendt-Tranholm, Rhea Sayani, Shreya Jagan, Afsheen Hossain, Jasmine Rosenfield, Andres Jesus Rodriguez Concepcion, Gregory Dussor, Armen N. Akopian, Theodore J. Price

## Abstract

Prolactin receptor (PRLR) isoforms, specifically the short (PRLR-S) and long (PRLR-L), are differentially expressed in pathological conditions and may be involved in the induction and maintenance of pain in females. PRLR sex-dependent actions are well studied in rodent models of chronic pain, but the translatability of these findings have only been assessed in electrophysiology studies that found consistent results with previous sex differences described in mice. To increase our understanding of the sexual dimorphic nature of prolactin actions on sensory neurons and our knowledge of how this body of work applies to the human nervous system, we characterized the expression of the short and long isoforms of PRLR with RNAscope *in situ* hybridization in human dorsal root ganglion (DRG) neurons. Based on data in mice, we hypothesized *PRLR-S* and *PRLR-L* would be equally expressed in human DRG neurons, with higher expression in females. To test this, we used DRG tissues recovered from male and female organ donors and cervix as a positive control. Our results demonstrated human DRG neurons express more *PRLR-L,* specifically transcript variant 6 (ΔS1), than *PRLR-S* in both sexes and that female donors had a higher expression of *PRLR* overall. In DRGs from female organ donors with a history of chronic pain we observed less *PRLR-L* expression, a finding consistent with previous studies in pain models in female mice. Expression for both *PRLR* isoforms in cervix was consistent with previous reports. An analysis of previously published RNA sequencing studies on human DRGs revealed that *PRLR* expression is confined mostly to *TRPV1-*positive nociceptors in humans, consistent with our RNAscope-based findings. We conclude that the *PRLR-L (ΔS1*) form is the major isoform expressed in human DRG neurons, that this expression is higher in females than in males, and that *PRLR-L (ΔS1)* expression is suppressed in chronic pain conditions in females.

## 1. Introduction

Prolactin (PRL) is a hormone produced mainly in the anterior pituitary gland that binds to the prolactin receptor (PRLR) and initiates downstream signaling cascades to regulate lactation, reproductive development, and cell differentiation and survival^1–5^. PRL and PRLR are well studied in the cancer field where an increase in PRL is known to upregulate cancer cell growth and survival in reproductive tissue by activating Janus Kinase 2/signal transducer and activator of transcription-5 (JAK2/STAT5) pathways^6–12^. The increase of PRL in serum is also linked to migraines^13–16^, hyperprolactinemia^17^, and other pain disorders in preclinical and clinical studies^18–21^. In rodent studies, PRL injections induce sensitization of dorsal root ganglion (DRG) sensory neurons in females that cause hyperalgesia, an effect that can be reversed with a PRLR antagonist^22^. Later studies identified that the short isoform variants of PRLR (PRLR-S, variants SF1a, SF1b) are responsible for these pronociceptive effects, while the long isoform (PRLR-L) is antinociceptive^23,24^. Pathological conditions, such as pain, increase circulating and/or locally produced PRL and alter the relative expression levels of PRLR-S and PRLR-L to favor pronociception in rodent models^24,25^. It is unknown whether this change in PRLR isoform expression also occurs in the human DRG (hDRG).

Humans have 16 protein-coding transcript variants (tv) for PRLR^26–30^. The most studied isoforms of PRLR are the long (tv1) and short (tv3,4) isoforms in the cancer and pain field. The canonical long form (tv1) includes the full-length of the PRLR membrane-bound receptor that is primarily responsible for mediating PRL signaling through Jak2 kinases and induce breast development and lactation^31,32^. The ΔS1 (tv6) variant contains a S1 motif deletion of exon 4 and 5 in the extracellular ligand-binding domain that causes a 7-fold reduced binding affinity for PRL compared to the canonical long form, but is otherwise functionally similar to the canonical long form and activates Jak2 to induce downstream pathways^29^. The short isoforms are alternatively spliced on exon 11 to produce SF1a (tv3) and SF1b (tv4)^30,33^. They are known to act as a dominant negative regulator and inhibit long form PRLR activity in breast cancer^34,35^ and to be upregulated and have pronociceptive effects in pain models in rodents^24^. The Δ7/11 (tv5) variant is a soluble prolactin-binding protein that lacks a transmembrane domain and inhibits PRL-induced cell proliferation by binding free PRL^36^.

The purpose of our study was to clarify our understanding of PRLR isoform expression in hDRG sensory neurons given the strong evidence for the functional divergence between isoforms in pathological conditions and the potential importance of this receptor in female-specific pain. Expression analysis was conducted using RNAscope *in situ* hybridization with specific probes against the long and short forms of PRLR in tissues procured from organ donors. Our findings provide insight into the expression of the PRLR isoforms in hDRG neurons which help inform the field of the therapeutic potential and limitations of PRLR as a target for pain.

## 2. Methods

### Tissue Procurement

DRGs were recovered from organ donors through a collaboration with the Southwest Transplant Alliance (STA) within 4 hours of cross-clamp from organ donors as previously described^37^. Upon removal from the body, DRGs were immediately frozen in powdered dry ice in the operating room, transported to UT Dallas on dry ice, and then stored at -80°C. Donor demographics, medical history, and samples utilized in each experiment can be found in Table 1. Three male and three female donors without chronic pain were used for the PRLR characterization experiments. As a result of troubleshooting the PRLR probes, a long-time frame occurred between experiments and different donors were used due to tissue availability. For the validation of the RNAscope *in situ* hybridization PRLR isoform probes, healthy (Cat # CB501526) and adenocarcinoma (Cat # CB538535) fresh frozen human endocervical tissue blocks were purchased from Origene and stored at -80°C.

**Table 1.** Human Donor Demographics.

| UTD ID | Sex | Age | Medical History | Experiment |
|---|---|---|---|---|
| UTD-DN0003 | M | 53 | DM, HLD, Hypertension, shoulder sx within last 6 months | Gen/L/S |
| UTD-DN0006 | M | 29 | Marijuana use, no other significant history | Gen/S |
| UTD-DN0091 | M | 22 | Drug use, DM2x3 year; 3 pack/yr smoking hx; | Gen/S |
| UTD-DN0147 | M | 19 | no significant history | L |
| UTD-DN0114 | M | 31 | Tylenol for occasional headaches, asthma as a child (resolved) | L |
| UTD-DN0001 | F | 34 | smoking and IV drug use | Gen/S |
| UTD-DN0042 | F | 44 | Depression, anxiety, headaches, IBS, sx: gall bladder removal, cyst removal from L breast | Gen/L/S |
| UTD-DN0122 | F | 19 | Drug use, tox all negative on testing, but alcohol | Gen/S |
| UTD-DN0166 | F | 38 | Asthma (albuterol); heart attack on this admission | L |
| UTD-DN0151 | F | 25 | on birth control; previous C-section | L |
| UTD-DN0261 | F | 53 | DM T1 since age 18; IBS; HTN; Heart Failure; Medications: DI, Fibro, HTN medication, **Hydrocodone for pain**, list incomplete; Allergy to sulfa drugs; Surgical hx: Wrist injury repair, Foot surgery for degenerative repair (5 yrs ago), Toe surgery (5 yrs ago), BL shoulder plates for degenerative repair (10 yrs ago), Embedded heart monitor, Port access placed in chest (15 yrs ago). | Pain L & S |
| UTD-DN0133 | F | 40 | Psoriatic Arthritis (methotrexate), Depression, Bipolar, **Oxy for Pain**, Clonopin for Anxiety, Unknown meds for ADHD, Birth Control, Narcolepsy, Thyroid, recent sterioids for Strep; PCN allergy; Sinus surgery June 2022, Asthma 10-15 yrs ago with inhaler use | Pain L & S |
| UTD-DN0106 | F | 48 | DM x 22 yrs, ESRD (dialysis), HTN x 22 yrs, CVA, Seizure disorder, Hepatic disfunction; Current Medications: Velafaxine, Furosemide, Propranolol, Trazodone, Vitamin D3, Nifedipine, Fluticasone, Novologinsulin pen (diabetes), Famotidine (reduce acid), Hydralazine (blood pressure), Sevelamer (dialysis), Mirtazapine, Lantus insulin (diabetes), Losartan; **Hydrocodone usage from 2012-2017 intermittantly**; Surgical history: Left foot partial amputation, Left thigh rod placement, Left knee surgery, Craniectomy, Cataract surgery both eyes. | Pain L & S |

All procedures were approved by the Institutional Review Boards at the University of Texas at Dallas. All human tissue procurement procedures and ethical regulations were approved by the Institutional Review Board at the University of Texas at Dallas under protocol Legacy-MR-15-237. Dorsal root ganglion (DRG) tissue was procured from organ donors through a collaboration with the Southwest Transplant Alliance, an organ procurement organization (OPO) in Texas. The Southwest Transplant Alliance obtains informed consent for research tissue donation from first-person consent (driver’s license or legally binding document) or from the donor’s legal next of kin. Policies for donor screening and consent are those established by the United Network for Organ Sharing (UNOS). OPOs follow the standards and procedures established by the US Centers for Disease Control (CDC) and are inspected biannually by the Department of Health and Human Services (DHHS). The distribution of donor medical information is in compliance with HIPAA regulations to protect donor privacy.

### Tissue Preparation

Frozen tissues were removed from the -80°C freezer and embedded in optimal cutting temperature (OCT) in a cryomold over dry ice to avoid thawing. Samples were then equilibrated to -20°C in the cryostat for 30 minutes before cryosectioning. Four 20 µM sections were mounted onto charged microscope slides for each donor. The slides were either immediately used for experiments or stored in the -80°C for up to one week.

### RNAscope *in situ* hybridization

The slides were fixed in cold 10% formalin for 15 minutes, followed by dehydration in 50%, 70%, and 100% ethanol (twice) for 5 minutes each at room temperature. The slides were briefly air dried and a hydrophobic barrier was drawn around each section using an ImmEdge PAP pen. The hydrophobic barrier was allowed to dry for 30 minutes before proceeding.

RNAscope *in situ* hybridization multiplex version 2 (manual assay, Advanced Cell Diagnostics (ACD) document number UM 323100) was conducted as instructed by ACD using the fresh frozen protocol. Hydrogen peroxide (ACD, Cat # 322381) was added to completely cover each section and incubated for 10 minutes at room temperature. Then the slides were washed twice in distilled water before proceeding to protease digestion. The protease reagent type and incubation time were optimized for each tissue as suggested by ACD. Human DRG was treated with protease III for 7-10 seconds, while human cervix received protease IV for an optimal 5 minutes before washing twice in 1x phosphate buffered saline (PBS, pH 7.4). The slides were singly covered in a probe mixture of channel 1 (C1), channel 2 (C2), and channel 3 (C3) probes at a 50:1:1 dilution, respectively, and placed in a prewarmed humidity control tray with wet filter paper. The control slides were similarly covered in either the positive or negative control probe. The humidity control tray was inserted into the HybEZ oven for 2 hours at 40°C to hybridize the probes. After hybridization, the slides were washed twice in 1x wash Buffer (ACD; Cat # 310091) before storage in 5X Saline Sodium Citrate buffer (Sigma; Cat # S6639) overnight at room temperature.

The next day, the slides were washed twice in 1X wash buffer before proceeding to AMP hybridization. All subsequent incubation steps were performed in the HybEZ oven at 40°C and followed by washing twice in 1x wash buffer. AMP 1 was added to cover the sections and incubated for 30 minutes, followed by AMP 2 for 30 minutes, and AMP 3 for 15 minutes. HRP-C1 reagent was added to the sections and incubated for 15 min. TSA Plus Akoya Dyes in Fluorescein, Cyanine-3, and Cyanine-5 (Akoya; NEL741001KT, NEL744001KT, NEL745001KT) were diluted in TSA multiplex buffer (ACD; Cat# 322809). Cyanine-3 and Cyanine-5 were diluted 1:1000, whereas Fluorescein was diluted 1:250. The appropriate akoya dye for labeling the C1 probe was added and incubated for 30 minutes. After washing, the HRP blocker (ACD; Cat # 323110) was applied for 15 minutes. The ensuing steps for HRP-C2 and HRP-C3 were identical to HRP-C1. The HRP-C2 and HRP-C3 steps were performed with their assigned akoya dye. After washing, DAPI (Cayman Chemical; Cat# 14285; 1:5000) was applied to each section for 5 minutes at room temperature and then washed in 1X PBS before air drying and cover-slipping (Globe Scientific; Cat # 1415-15) each slide with Prolong Gold Antifade mounting medium (Fisher Scientific; Cat # P36930). The slides were cured in the dark at room temperature overnight, sealed with nail polish, and stored in 4°C.

RNAscope Probes against *PRLR* with preferential detection of specific transcript variants were designed by ACD as described in **Table 2**. *PRLR* probes were paired with the nociceptor marker, *TRPV1* (transient receptor potential cation channel subfamily V member 1). For human cervix experiments, probes against *MKNK2 (*MAPK Interacting Serine/Threonine Kinase 2) and *DCN* (Decorin) were used alongside the *PRLR* probes as a positive control. A third channel was left unstained in order to visualize background lipofuscin. Probes against housekeeping mRNAs (*UBC*, *PPIB*, *POLR2A*; ACD Cat # 320861) were used as a positive control on a single section from each donor. A probe against the bacterial DapB mRNA (ACD; Cat # 320871) was used as the negative control.

**Table 2.** RNAscope *in situ* Hybridization probes.

| Probe | Type | Catalog # | Lot# | Variants detected | Detected BP | Probe |
|---|---|---|---|---|---|---|
| Hs-PRLR-C2 | PRLR-General | 542551-C2 | 21273C | **tv1 (long)**, tv6 (ΔS1), tv2, tv3 (SF1a), tv4 (SF1b) | 222-1271 | 20ZZ |
| Hs-PRLR-tv1tv6-O1-C1 | PRLR-L | 466731-C1 | 23219B | tv1 (long), **tv6 (ΔS1)** | 1336-2687 | 20ZZ |
| Hs-PRLR-O1-C2 | Custom PRLR-S | 1806011-C2 | 24366D | tv3 (SF1a), tv4 (SF1b), tv5 (Δ7/11), tv7 (non-coding) | 1103-1772 | 11Z |
| Hs-TRPV1-C1 and C2 | TRPV1 | 415381 | 23332A, 23108A | NA | 1922-2905 | 20ZZ |
| Hs-MKNK2-C1 | MNKN2 | 855491-C1 | 20090B | NA | 239-1636 | 20ZZ |
| Hs-DCN-C3 | DCN | 589521-C3 |  | NA | 171-1225 | 20ZZ |

A long delay occurred between the non-pain and female chronic pain donor PRLR characterization experiments. During this time frame, RNAscope *in situ* hybridization was optimized to reliably detect up to 4 channels instead of 2. This allowed the detection of *PRLR-L*, *PRLR-S*, and *TRPV1* simultaneously with an empty channel for autofluorescence in the female chronic pain donors.

### Imaging and Analysis

An Olympus FV3000 or FV4000 confocal microscope was utilized to capture the RNAscope *in situ* hybridization images. The images were taken at x20 magnification for DRG neurons. Three images were acquired for each DRG tissue section, and 3-4 sections were imaged per donor. The raw DRG image files were adjusted for brightness in the Olympus CellSens software and each neuron was assessed for the presence of *PRLR* and/or the nociceptor marker, *TRPV1*. The *PRLR-S* and *PRLR-L* RNAscope experiments were performed separately in order for there to be a channel for *TRPV1* labeling and a channel for lipofuscin background to be imaged. The average number of mRNA puncta was counted for each channel for the negative control and averaged to determine the threshold for each channel. Neurons were considered positive for *PRLR* if they contained 5 or more puncta. Neurons were considered positive for *TRPV1* if they contained 15 or more puncta. The neuron diameter was measured using the polyline tool in CellSens. The female chronic pain RNAscope experiment was imaged on the Olympus FV4000 with *PRLR-S, PRLR-L, TRPV1,* DAPI, and an blank channel for autofluorescence. Puncta that auto-fluoresced in each channel was considered background lipofuscin and not regarded as target-specific signal. The puncta threshold remained the same as the experiments performed in non-pain donors.

In cervical tissue, an Olympus FV4000 was used to capture three 40X z-stack images from 2-3 sections. Probes against *PRLR* isoforms with *MKNK2* or *DCN* as positive controls were used with DAPI and a blank channel for background autofluorescence. A qualitative assessment was performed to confirm PRLR detection with RNAscope because PRLR is known to be enriched in female reproductive organs. Puncta detected in the blank channel, and as autofluorescence in other channels, was considered background autofluorescence and not target specific signal.

## Statistical Analyses

GraphPad Prism 10.0.2 (GraphPad, La Jolla, CA) was used for all statistical analyses of data. Two-way ANOVA with Bonferroni’s *post-hoc* test was used to calculate the statistical significance between groups when appropriate. Data are provided as the mean ± SEM. Differences were deemed to be statistically significant when p < 0.05.

## 3. Results

For this study, we used *PRLR-General*, *PRLR-L*, and custom made *PRLR-S* RNAscope probes from ACD. The *PRLR-General* probe binds exons 3-8 in the extracellular domain and detects transcript variants (tv) 1, 2, 3, 4, 6 at varying sensitivities due to the lack of exon 4 and 5 in tv6 (Table 2, Fig 1). The *PRLR-L* probe binds exon 10 and detects tv 1 and 6, with a higher sensitivity for tv6 than the *PRLR-General* probe (Table 2, Fig 1). The custom *PRLR-S* probe binds exon 11 and detects tv 3, 4, 5, and 7 (non-coding mRNA) (Table 2, Fig 1). We expected the *PRLR-General probe* to bind both long and short isoforms and be co-expressed with the *PRLR-L* and *PRLR-S* probes.

**Figure 1.**
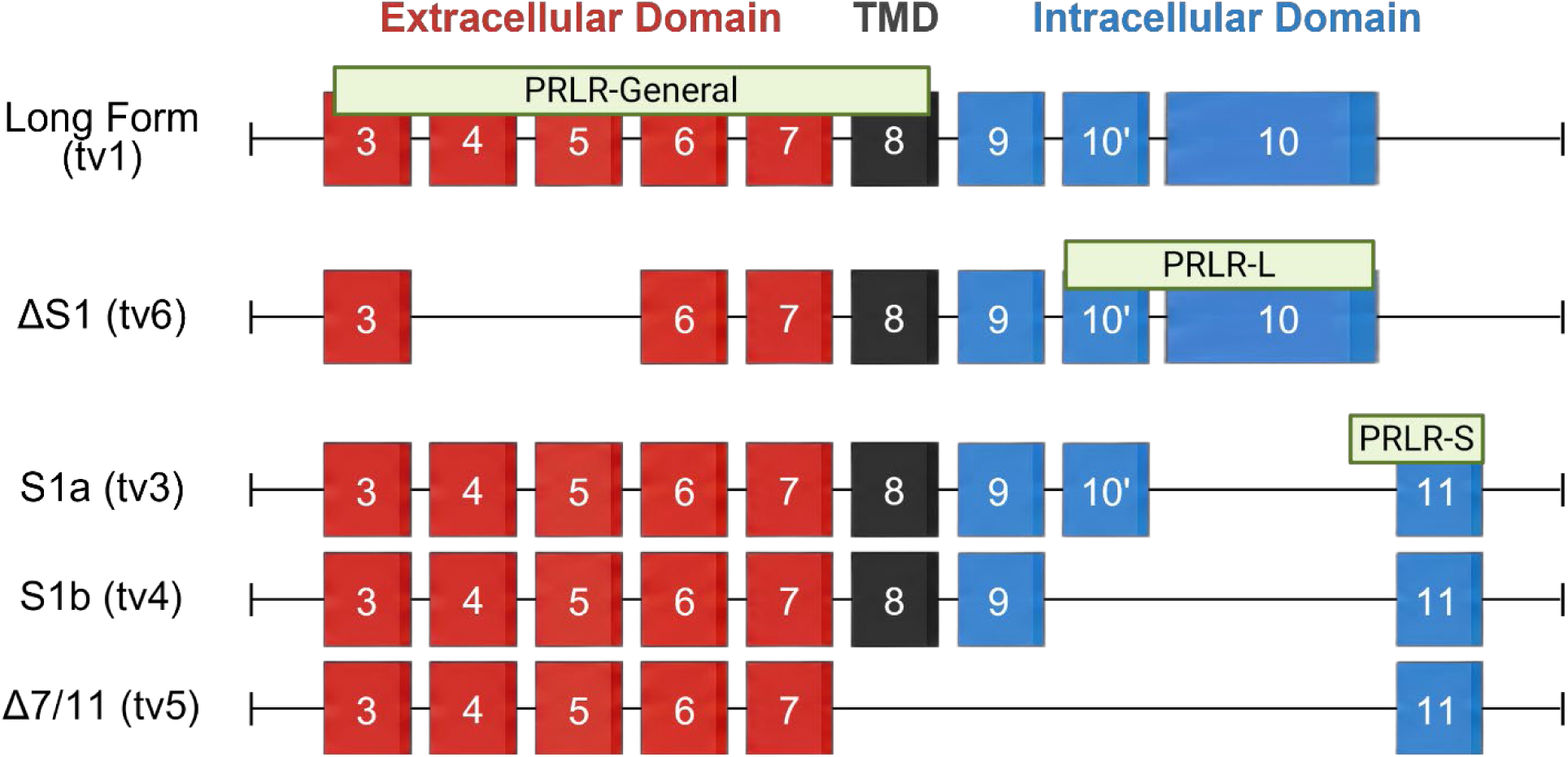
Schematic representation of human PRLR isoforms and the binding location of PRLR *in situ* hybridization probes. TMD, transmembrane domain. Adapted from reference^28^.

To validate the *PRLR* RNAscope probes, we used human endocervical tissue because bulk RNA- sequencing data demonstrated abundant *PRLR* mRNA (20.1-37.09 nTPM) expression in this tissue^38^. There are conflicting reports on whether the long or short form of *PRLR* is more abundantly expressed in normal or cancerous tissue. Previous immunohistochemical studies reported low PRLR protein levels in normal cervix tissue, but a significant upregulation of PRLR levels in cervical cancer tissue^39^. The same group also showed *PRLR-L* as the dominant isoform expressed in cervical cancer. However, other studies have shown the upregulation of PRLR-S over PRLR-L in tumor formation^40^ and cancer development^41^. Hematoxylin and eosin staining was performed and revealed glandular or columnar epithelial cells, indicating the tissue was from the endocervix^42^ (Fig 2A). RNAscope *in situ* hybridization (ISH) with *PRLR-L*, *PRLR-S*, and *PRLR- General* were performed on healthy and adenocarcinoma endocervical tissue. Bulk RNA- sequencing data indicated *MKNK2* and *DCN* had high mRNA expression in cervical tissue^38^ and were used as additional positive controls. The results demonstrated that *PRLR-L* (see Table 2 for summary of probes and transcript variants) was expressed more abundantly in healthy endocervical tissue, whereas *PRLR-S* and *PRLR-General* were more robust in the adenocarcinoma endocervix (Fig 2B, C). Although both the *PRLR-L* and *PRLR-General* probes detect long form (tv1) and ΔS1 (tv6), the *PRLR-L* probe is more sensitive to ΔS1 (tv6) (Fig 1). The expression of the *PRLR* probes was amassed in the stromal cells of the endocervical tissue. In support of previous studies, our results demonstrated an increase in long form, SF1a, and SF1b in cancerous cervical tissue.

**Figure 2.**
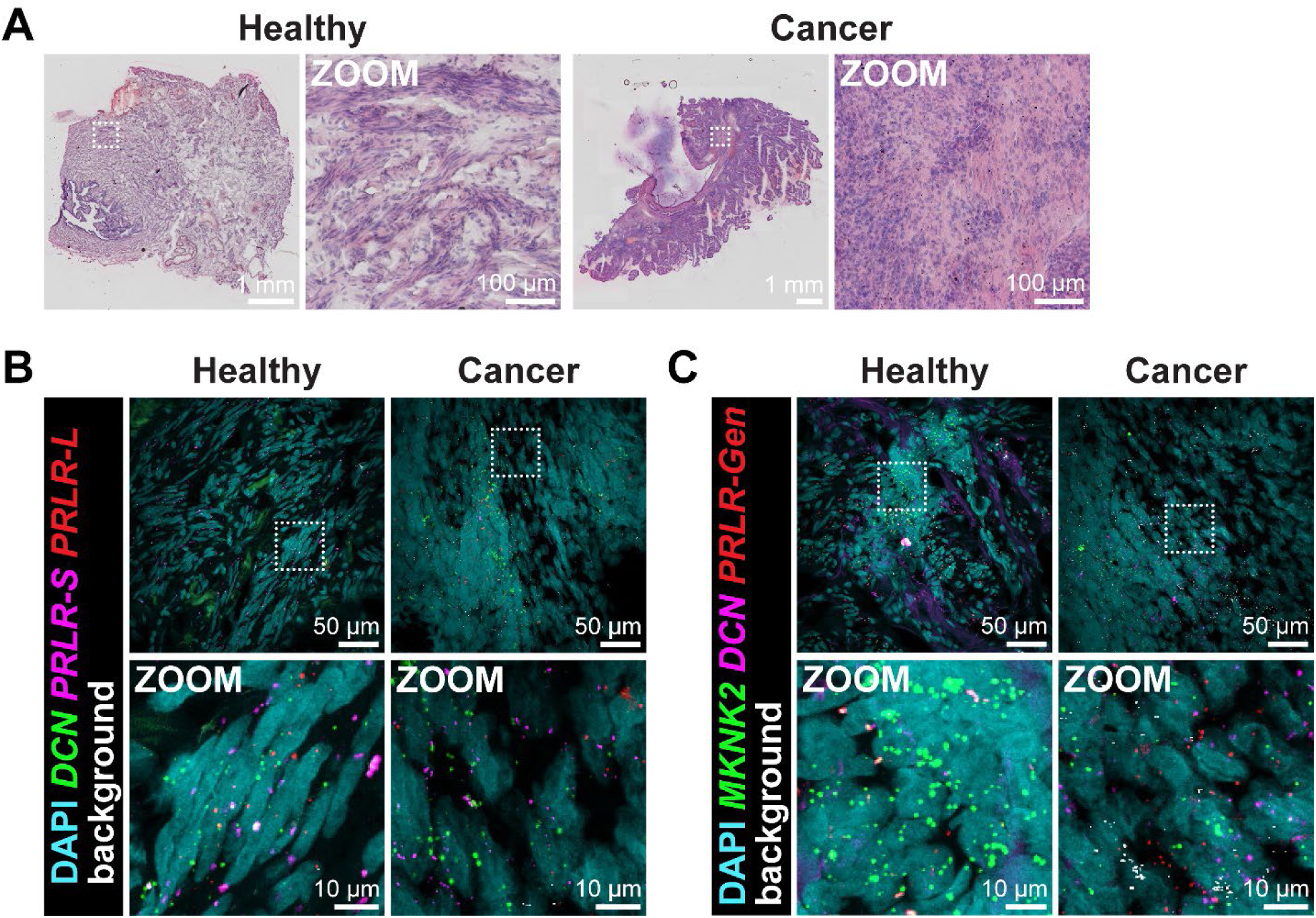
Validation of *PRLR* probes in human endocervical tissue. **A)** Representative images of hematoxylin and eosin stain of endocervical tissue from a healthy and adenocarcinoma donor. Inset displays digital zoom images of stromal cells. **B)** Representative 40X images of human endocervical tissue labeled with RNAscope *in situ* hybridization for DAPI (cyan), *DCN* (green), *PRLR-S* (magenta), *PRLR-L* (red), and background lipofuscin (white). **C)** Representative 40X images of human endocervical tissue labeled with RNAscope ISH for DAPI (cyan), *MKNK2* (green), *DCN* (magenta), *PRLR-General* (red), and background lipofuscin (white).

Once we validated the *PRLR* RNAscope ISH probes, we tested them on male and female DRG tissues. We detected very little *PRLR-General* in hDRG neurons (Fig 3A), despite our bulk sequencing data demonstrating its presence in mouse and human DRG. The probe design of the *PRLR-General* probe should detect both short and long isoforms (Fig 1). Male and female donors demonstrated a low level of *PRLR-General+/TRPV1-* expression in their DRG neurons at 0.19% and 1.29%, respectively. Male donors had twice the amount of *PRLR-General+*/*TRPV1+* co- expressing neurons than female donors at 4.56% and 2.11%, respectively (Fig 3B). In both histograms, it was observed that smaller diameter neurons (<50µm) were *TRPV1+* only and larger neurons (>60µm) were more likely to be *PRLR-General+/TRPV1-* or co-expressing *PRLR- General+*/*TRPV1+* neurons. Collectively, these results demonstrate *PRLR-General* is lowly expressed overall in human DRG and there are no significant differences in expression between male and female donors.

**Figure 3.**
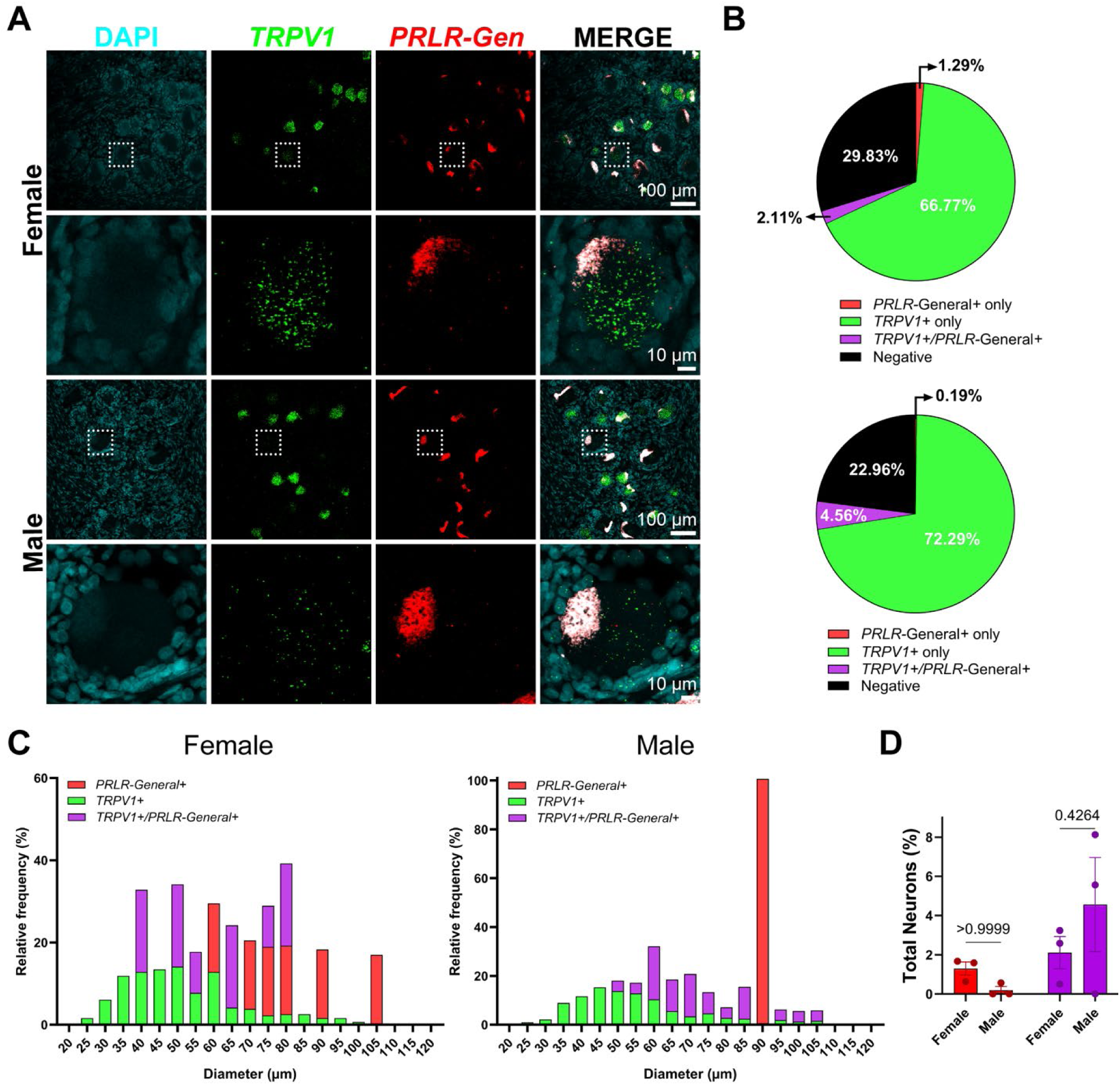
*PRLR-general* mRNA is lowly expressed in human DRG neurons. **A)** Representative 20X images of human female and male DRG labeled with RNAscope *in situ* hybridization for DAPI (cyan), *TRPV1* (green), and *PRLR-general* (red). Inset shows digital zoom of a single neuron with expression of *TRPV1 and no PRLR-general expression*. **B)** Pie chart representation of total neuron subpopulations in human DRG between female and male donors. **C)** Histogram displaying the size profile of all neuronal subpopulations in female and male human DRG. **D)** Comparison of neuronal subpopulations demonstrate there are no significant differences between sex. Two-way ANOVA with Bonferroni *post hoc*.

The *PRLR-L* and a custom made *PRLR-S* probes were obtained to further delineate isoform expression. We observed the *PRLR-L* probe robustly detected *PRLR* signal compared to the *PRLR-General* probe in hDRG neurons (Fig 4). The *PRLR-L* probe also detects long form and ΔS1, but with a higher sensitivity for tv6 than the *PRLR-General* probe. The expression of *PRLR- L* was more robust (Fig 3A) in the DRG neurons of both male and female donor DRG neurons compared with *PRLR-General*. This suggests that ΔS1 may be the dominant long isoform in hDRG neurons.

**Figure 4.**
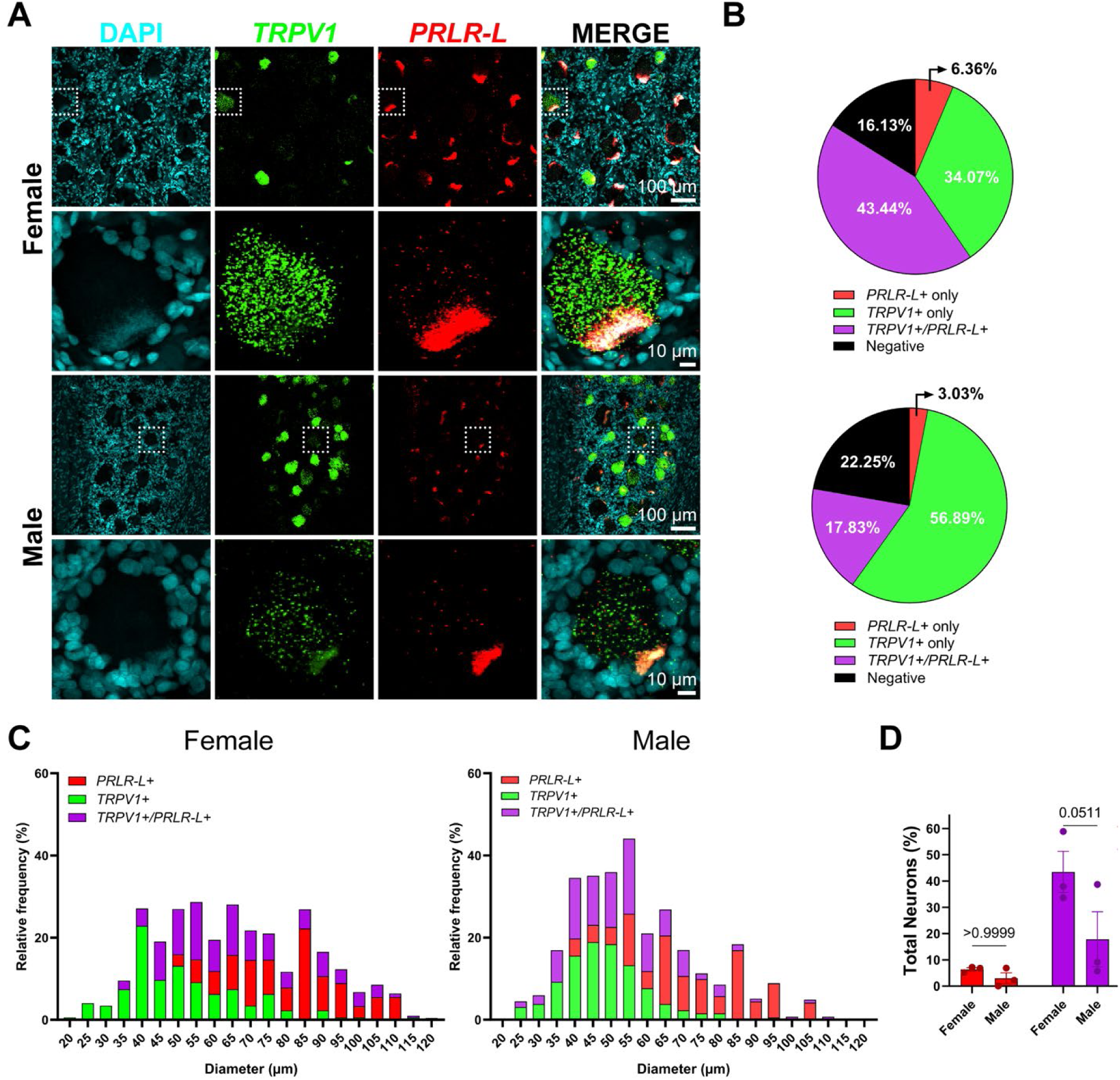
*PRLR-L* mRNA is expressed more robustly in female human DRG neurons. **A)** Representative 20X images of human female and male DRG labeled with RNAscope *in situ* hybridization for DAPI (cyan), *TRPV1* (green), and *PRLR-L* (red). Inset shows digital zoom of a single neuron with co-expression of *TRPV1* and *PRLR-L*. **B)** Pie chart representation of total neuron subpopulations in human DRG between female and male donors. **C)** Histogram displaying the size profile of all neuronal subpopulations in female and male human DRG. D) Comparison of neuronal subpopulations demonstrate a trending significant difference between PRLR+/TRPV1+ neurons between male and females. Two-way ANOVA with Bonferroni *post hoc*.

Female DRG neurons expressed twice the percentage of *PRLR-L+/TRPV1-* (6.36%) and *PRLR- L+/TRPV1+* (43.44%) than male DRGs (3.03%, 17.83%) (Fig 3B). We again observed that small diameter neurons (<50µm) were mostly *TRPV1+* neurons, whereas *PRLR-L*+ neurons were mostly in larger diameter neurons (>60µm) in both male and female DRG neurons (Fig 4C). Interestingly, the male histogram was shifted to the left, whereas the female histogram was more normally distributed. The number of neurons showing expression of *PRLR-L+/TRPV1+* were on the border of statistically significant (p=.051) between males and females (Fig 4D).

The *PRLR-S* probe detects both short form variants (SF1a, SF1b), as well as the prolactin-binding protein Δ7/11, and a non-coding RNA variant. *PRLR-S* was lowly expressed in the DRG neurons of both male and female donors (Fig 5A). Females expressed *PRLR-S+* in only 1.94% of DRG neurons, whereas males expressed it in only 0.21% (Fig 5B). We observed that *PRLR-S*+ neurons were expressed in small to large neurons in female DRG’s, but exclusively in *TRPV1+* large diameter neurons in males (Fig 5C). There were no significant sex differences in neuronal subpopulation expression (Fig 5D).

**Figure 5.**
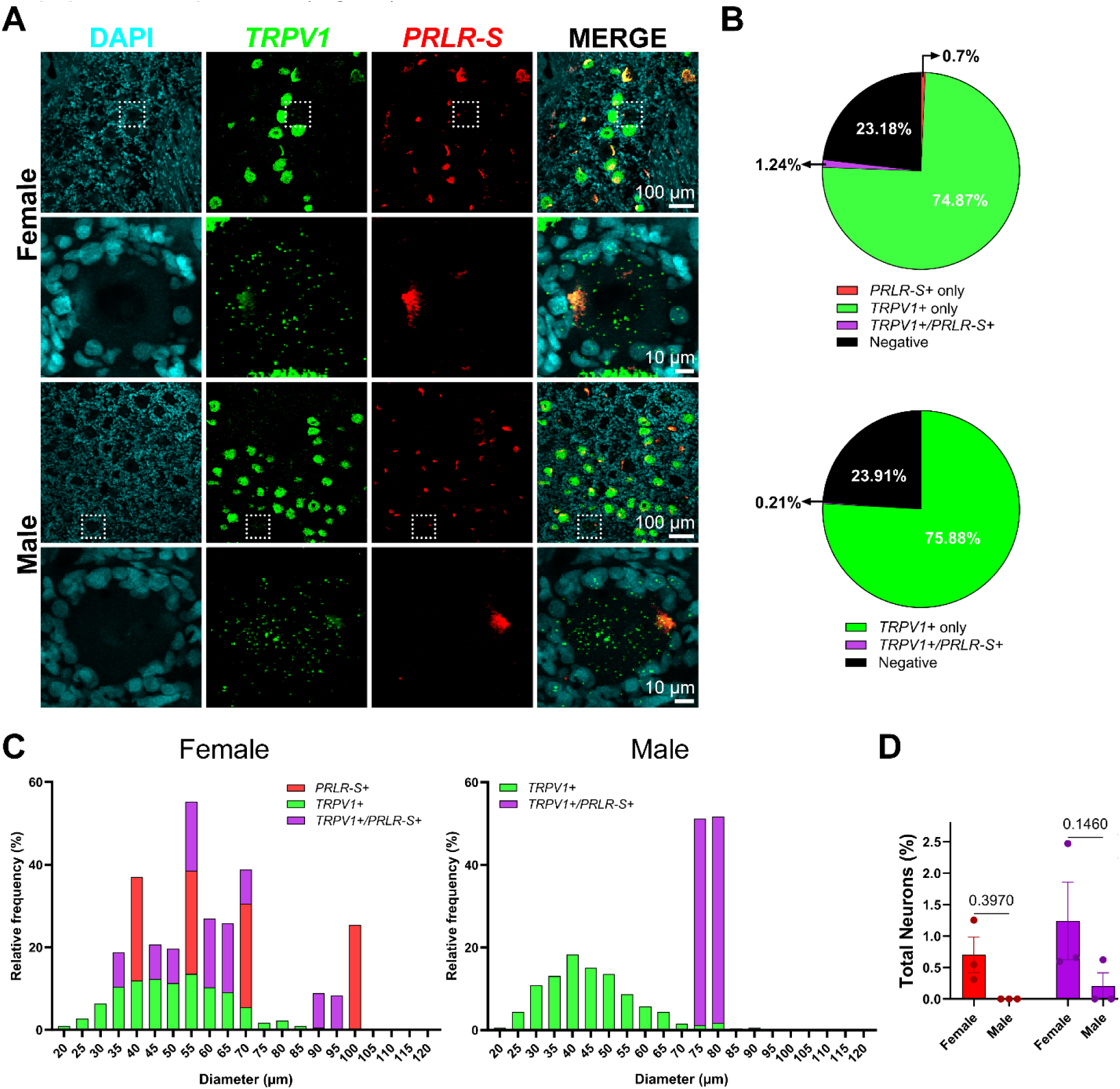
*PRLR-S* mRNA is lowly expressed in human DRG neurons. **A)** Representative 20X images of human female and male DRG labeled with RNAscope *in situ* hybridization for DAPI (cyan), *TRPV1* (green), and *PRLR-S* (red). Inset shows digital zoom of a single neuron with expression of *TRPV1* only. **B)** Pie chart representation of total neuron subpopulations in human DRG between female and male donors. **C)** Histogram displaying the size profile of all neuronal subpopulations in female and male human DRG. **D)** Comparison of neuronal subpopulations demonstrate there are no significant differences between sex. Two-way ANOVA with Bonferroni *post hoc*.

A recent study in rodents found that a hind paw incision significantly decreased PRLR-L protein levels in mouse DRG but had no change in PRLR-S ^25^. The authors also observed the prolactin neutralizing antibody could significantly decrease human and mouse DRG neuron excitability in prolactin treated neurons as well as attenuate pain behavior. Thus, we asked whether pain conditions in humans could also lead to a decrease in *PRLR-L* expression. We performed RNAscope *in situ* hybridization with *PRLR-S*, *PRLR-L*, and *TRPV1* probes in female donors with pain conditions and found significantly more *PRLR-L* mRNA expression in females without pain (Fig 6A-D). This finding corroborates studies implicating the long form as protective against pain ^23,24^. We did not observe an increase in *PRLR-General* in the female pain donors (supplemental Fig 1), thus the analysis was not performed. The female pain donors demonstrate the vast majority of their *TRPV1+* large diameter neurons (> 60 μm) co-expressed *PRLR+*, with the largest neurons (>100 μm) exclusively expressing *TRPV1+/PRLR-S+*. We also compared the percentage of total neuronal populations between male and female DRG neurons (Fig 6D). The data shows there are more female neurons co-expressing *PRLR-L+/TRPV1+* than any other population in either sex. This aligns with previous work demonstrating female rodents have more functional PRLR that drives an effect on nocifensive behavior and influences neuronal hyperexcitability in response to PRL^22,43,44^.

**Figure 6.**
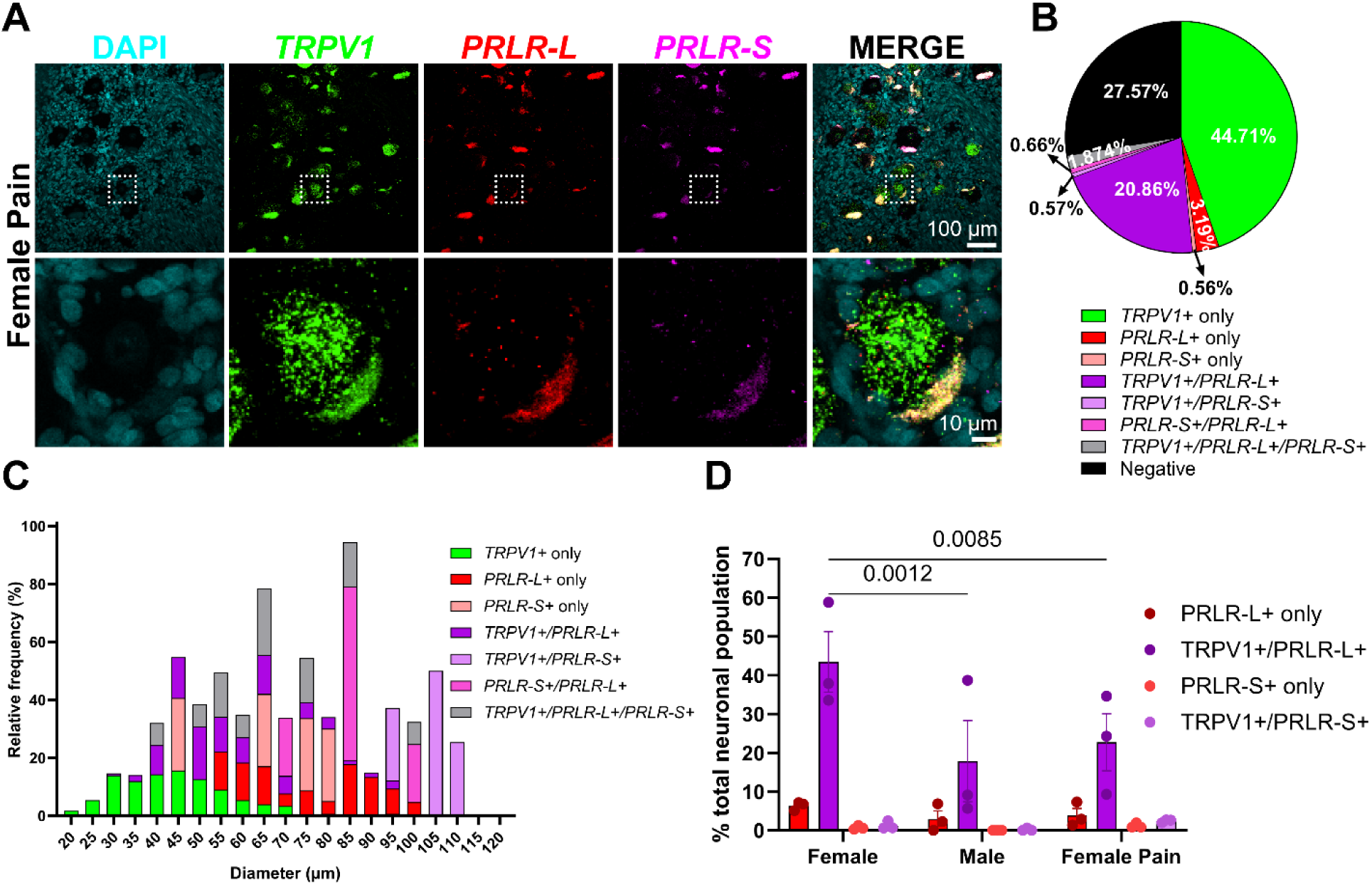
*PRLR-L* mRNA is reduced in females with chronic pain. **A)** Representative 20X images of human female DRG labeled with RNAscope *in situ* hybridization for DAPI (cyan), *TRPV1* (green), *PRLR-L* (red), and *PRLR-S* (magenta). Inset shows digital zoom of a single neuron with expression of *TRPV1 and PRLR- L*. **B)** Pie chart representation of total neuron subpopulations in female hDRG from chronic pain donors. **C)** Histogram displaying the size profile of all neuronal subpopulations in female chronic pain hDRG. **D)** *TRPV1+/PRLR-L+* was expressed more robustly in females without pain than those with pain. Two-way ANOVA with Bonferroni *post hoc*.

## 4. Discussion

In this study, we investigated the mRNA expression of the *PRLR-General*, *PRLR-S*, and *PRLR-L* probes in hDRG tissue. We hypothesized *PRLR-L* and *PRLR-S* would be expressed equally in hDRG neurons, and that female donors would have more *PRLR* expression overall. We also hypothesized that chronic pain donors would have upregulated *PRLR-S* or reduced *PRLR-L* in DRG neurons compared with non-pain donors. Our results partially supported the hypothesis by demonstrating that females had more *PRLR-L* expression than males, but only low amounts of *PRLR-S* expression in both sexes. Surprisingly, we identified *PRLR-L* ΔS1 (tv6) as the likely dominant long isoform in human DRG neurons instead of the canonical long form (tv1). Finally, we found that female chronic pain donors exhibited reduced *PRLR-L* ΔS1 compared to non-pain female donors, but no significant change in *PRLR-S* expression. Altogether our main findings were that 1) *PRLR-L ΔS1* may be the dominant isoform in human DRG neurons, 2) females express more *PRLR-L* ΔS1 in *TRPV1*+ neurons than males, 3) human DRG neurons express low levels of *PRLR-S*, and 4) *PRLR-L* ΔS1 expression is reduced in female chronic pain donors.

Our results indicate that *PRLR-L ΔS1* may be the dominant isoform in human DRG neurons rather than the canonical *PRLR-L* (Fig 6). This contrasts with cancer studies in breast and cervical tissues where the canonical PRLR-L is dominant^31,39,45^. Although it is possible that PRLR-L ΔS1 was not delineated from the canonical PRLR-L in these studies. The full functional role of PRLR- L ΔS1 is unclear as the exclusion of exon 4-5 results in a truncated extracellular binding domain that impacts PRL binding. However previous results indicate limited functional impact based on the ability of PRLR-L ΔS1 to activate downstream pathways through JAK2^29,32,45–49^. PRLR-L ΔS1 has a 7-fold reduction in binding affinity to PRL than the canonical PRLR-L and does not self- antagonize at high concentrations of PRL^50,51^. Presumably, this would allow PRLR ΔS1 to continuously signal downstream pathways and sensitize DRG sensory neurons. Potential explanations for PRLR-L ΔS1 as the dominant isoform in the hDRG could be that various tissues and species have diverse expression levels of the PRLR isoforms. One study found differentially expressed *PRLR-L ΔS1* and *PRLR-L* in human tissues, specifically they found higher *PRLR-L ΔS1* mRNA expression in human placenta, kidney, and uterus compared to the canonical *PRLR- L*^29^. Their result aligned with another group where they identified absent *PRLR-L* mRNA expression in healthy cervical tissues but an upregulation of the *PRLR-L* in cervical cancer samples^39^. These findings coincide with our own observation of upregulated canonical *PRLR-L* and downregulated *PRLR-L ΔS1* in adenocarcinoma endocervical tissue (Fig 2). In contrast, mouse DRG are reported to either express similar levels of PRLR-S and PRLR-L across sexes or an upregulation in females^24,44^. To our knowledge there have been no investigations on the PRLR ΔS1 isoform in rodent or human DRG neurons.

We expected females to have more *PRLR-L* expression than males due to the female-specific role of PRL and PRLR in previous literature and our own studies^22,24,43,52^. Our results did not support the hypothesis that *PRLR* isoforms would be equally expressed, we found more *PRLR-L ΔS1* expression than other isoforms in both sexes. This result is in contrast with multiple reports in rodent DRG neurons ^24,44,53^. These results could be attributed to gonadal hormone levels, age, or species differences. Estrogen is known to upregulate only *PRLR-L* expression in rodent trigeminal neurons^52^ and reproductive tissues^22,24,54,55^, which could explain the sex difference in *PRLR-L ΔS1* expression. In the same study, acute PRL also sensitized TRPV1 capsaicin responses^52^. Given our finding that PRLR-L ΔS1 is the dominant isoform in the hDRG, it may suggest that PRLR-L ΔS1 is involved in the sensitization of DRG neurons. Future functional studies are needed to delineate the PRLR isoforms involved in DRG neuron sensitization and understand the mechanisms by which PRLR-L ΔS1 sensitizes DRG neurons.

Most pain studies attribute the PRLR-S with pronociception and PRLR-L with antinociception in the trigeminal ganglion (TG) and DRG sensory neurons in rodents^23–25^. These studies were all performed in rodents and present translatability challenges, especially since we demonstrate that less than 2% of human DRG neurons express *PRLR-S*. A limitation of previous investigations is the lack of specific antibodies and probes to detect PRLR isoforms and the isolation of neuronal expression. For instance, it is unknown whether the PRLR-L antibody also detected the intermediate form or if the PRLR-S antibody detected Δ7/11 or other truncated variants. The quantification of PRLR isoform expression in rodent DRG tissue using western blot or RT-PCR includes nerve fibers and glia cells and thus is not specific to neuronal expression of PRLR^22,44^. Recently, papers from the Porreca group reported PRLR protein expression exclusively in cultured female donor DRG neurons and overnight incubation with high dose hPRL increased their excitability only in female donors^25,56^. Their results supplement our human findings by demonstrating a sex difference in PRLR expression in hDRG neurons and demonstrating PRL’s ability to induce increased excitability in human sensory neurons. The authors assumed that since PRLR-S induced nociception in rodents, the excitability of hDRG neurons was also due to PRLR-S. Our results demonstrate that PRLR-L ΔS1 is present in ∼50% of DRG neurons in women and PRLR-S in ∼2%, thus making it unlikely that PRLR-S is involved in nociception. While it is plausible that our probe did not detect short forms, or the short form mRNA does not accurately reflect protein expression, this is unlikely as our PRLR-S probe detected the short forms in adenocarcinoma cervical tissue and it is known to have a longer half-life than PRLR-L^40,57^. We favor the idea that the PRLR-L ΔS1 is more likely to be involved in hDRG neuron excitability and nociception, not PRLR-S. The PRLR-L ΔS1 has a 7-fold lower binding affinity than the canonical long form and does not self-antagonize at higher concentrations of PRL^29^. Stratton et al used a high concentration of PRL at 50nM overnight, whereas others have used lower concentrations and shorter treatment times (5-20 minutes)^22,53,58^ on rodent DRG neurons. The high concentration and long treatment time of PRL could activate more long-term pathways that sensitize DRG sensory neurons. Prior research found that acute PRL treatment sensitized neurons through TRPV1 and other voltage-gated channels and long term PRLR-L signaling is more involved in JAK/STAT5 and MAPK pathways that have transcription activation but may have the potential for rapid signaling through PKC^58^. The PRLR-L may also induce the sensitization of DRG neurons by activating JAK2/STAT3 to drive the transcription of pro-inflammatory cytokines^59–61^. Future studies will determine *PRLR-L* ΔS1 downstream signaling in hDRG neurons.

Our results in female pain donors demonstrate a significant reduction in *PRLR-L* ΔS1 in TRPV1+ neurons from ∼45% to ∼20% (Fig 3 & 6). There are two likely reasons for the reduction in *PRLR- L* ΔS1: 1) *PRLR-L ΔS1* could be protective against pain in the DRG of a healthy person and chronic pain could lead to the loss of this protective mechanism or 2) *PRLR-L ΔS1* signaling is involved in nociception and chronic pain may downregulate *PRLR-L ΔS1* in sensory neurons to suppress hypersensitivity. There are many studies supporting the first hypothesis^22–24^, but it does not explain the low level of *PRLR-S* found in the hDRG. The second hypothesis is less supported in rodent literature but would be more likely due to the absence of *PRLR-S* in the human DRG. If PRLR-L ΔS1 is the dominant isoform expressed in the hDRG, then it must be involved in nociception as demonstrated by clinical studies demonstrating the pronociceptive role of PRL^14,24,55,62–64^.

In conclusion, our results utilizing human DRG tissue diverge from rodent studies. We reveal *PRLR-L ΔS1* may be the dominant isoform that induces pronociceptive signaling in human DRG neurons and that the expression of this isoform is reduced in female chronic pain donors. Future studies to delineate the protein expression and signaling properties of the PRLR isoforms in human DRG neurons are needed to understand the therapeutic impact of PRLR for pain in humans.

