## Supplementary material for "Prolactin Receptor Long Isoform ΔS1 Expression is Reduced in the Dorsal Root Ganglia of Humans with Chronic Pain"

**This PDF file includes:**

Figure S1

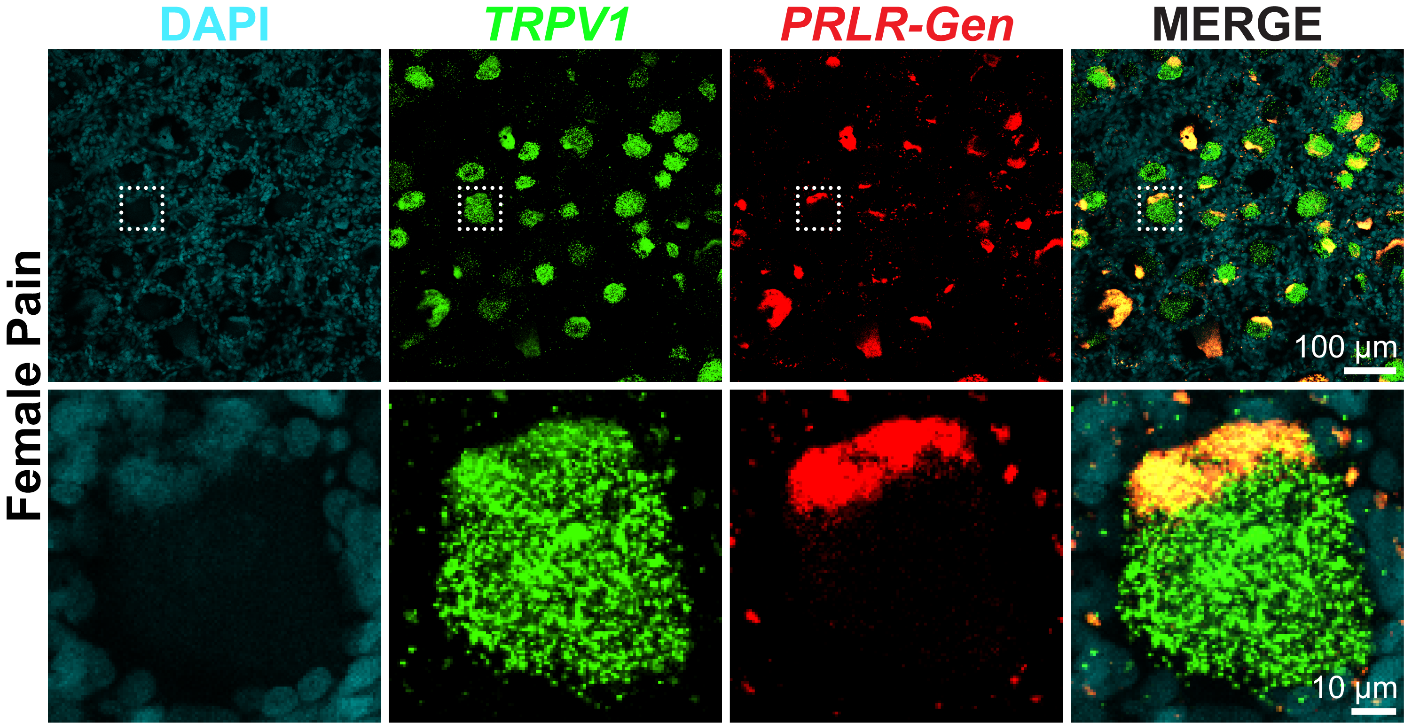

**Figure S1.** Representative human DRG RNAscope *in situ* hybridization for DAPI (cyan), *TRPV1* (green), and *PRLR-General* (red) in a female chronic pain donor. Digital zoom images show a single neuron with *TRPV1* expression but an absence of *PRLR-General* mRNA expression.
